# Single-cell characterization of tumor immune landscapes in colorectal cancer humanized mice

**DOI:** 10.64898/2026.06.01.729295

**Authors:** Sanne Bootsma, Job Saris, Andrew Y.F. Li Yim, Kristiaan J. Lenos, Felipe A. Vieira Braga, Joep Grootjans, Louis Vermeulen

## Abstract

Colorectal cancer (CRC) displays inter-patient heterogeneity in molecular tumor features and immune cell composition, which influence therapy response. Humanized immune system (HIS) mouse models offer a promising *in vivo* model to study human tumor-immune interactions, yet their ability to recapitulate the CRC tumor immune microenvironment at single-cell resolution remains incompletely defined. Here, we performed single-cell RNA sequencing of systemic and tumor-infiltrating human immune cells in HIS mice bearing human CRC tumors and benchmarked these data against reference datasets of healthy human spleens and primary CRC tumors. Major immune lineages and transcriptional programs characteristic of the human systemic immune compartment were identified, and HIS mouse tumors developed complex, human-like immune infiltrates. Tumor-infiltrating immune cells comprised diverse T cell, myeloid, natural killer, and B cell populations, including exhausted T cell states marked by expression of *PDCD1, TIGIT, HAVCR2, LAG3*, and *CTLA4*. We further demonstrate CRC consensus molecular subtype-associated spatial differences in immune infiltration. Collectively, our findings support the use of HIS mice as a relevant model for studying CRC immune landscapes and preclinical evaluation of immunomodulatory therapies.

## INTRODUCTION

Colorectal cancer (CRC) is a heterogeneous disease, with substantial inter-patient variability in molecular tumor characteristics and therapeutic responses. Cancer immunotherapy with immune checkpoint blockade (ICB) has revolutionized the treatment of microsatellite unstable (MSI) CRC; however, these tumors account for only approximately 15% of all CRC cases^1-3^. The majority of microsatellite stable (MSS) CRCs are considered immunologically cold and only show very limited clinical responses to ICB. A better understanding of the interactions between tumor cells and immune cells is required for adequate patient selection and the development of new immunomodulatory treatments effective in the MSS subgroup. This ultimately requires *in vivo* models that mimic both the complexity of human cancer cells and the human immune system.

To study the tumor-immune interactions *in vivo*, genetically engineered or syngeneic mouse cancer models are typically used. However, these models often fail to fully recapitulate human tumor-immune interactions^4^. To circumvent this limitation, humanized immune system (HIS) mice have been developed. In these models, human hematopoietic stem cells (HSCs) are injected into immunodeficient mice, resulting in partial reconstitution of a human immune system. Several studies have shown successful engraftment of human patient-derived xenografts (PDX) in HIS mice, with variable tumor immune cell infiltration and sensitivity to ICB between different CRC PDX models^5-8^. This variability suggests that tumor-intrinsic features may shape the composition and functional state of the tumor immune microenvironment (TIME) and underscores the need for detailed immune profiling of these models.

To capture the heterogeneity and complexity of CRC, the gene expression-based consensus molecular subtype (CMS) classification system, as well as several related taxonomies, have been developed^9-11^. The CMS subtypes differ in their biological characteristics and clinical behavior and have been reported to show distinct immune features in patients. Specifically, CMS1 tumors represent MSI CRC with high immune infiltration and abundant cytotoxic lymphocytes, CMS2 and CMS3 are largely non-immunogenic, and CMS4 tumors display an inflamed yet immunosuppressive immune signature that is associated with poor prognosis^12^.

In this study, we aimed to systematically assess the extent to which HIS mice recapitulate the human systemic and tumor immune landscapes of CRC at single-cell resolution. We performed single-cell RNA sequencing (scRNA-seq) of human immune cells from both the systemic immune compartment and subcutaneous human CRC tumors grown in HIS mice. To enable controlled tumor-intrinsic stratification, we employed CMS-classified human CRC cell line models. We compared our data with publicly available scRNA-seq datasets of healthy human spleens^13^ and primary human CRC tumors^14^, thereby benchmarking immune cell composition and transcriptional programs. In particular, we evaluated the presence of immunologically relevant T cell states, including features of T cell exhaustion that are central to the mechanism of action of ICB therapy and are largely reproduced in HIS mice, supporting the utility of this platform for mechanistic studies and preclinical evaluation of immunomodulatory therapies in CRC.

## MATERIALS & METHODS

### Cell culture

Cell line Hutu80 (CMS4) was cultured in Dulbecco’s modified Eagle’s medium/F-12 (DMEM/F12) medium with L-glutamine, 15 mM HEPES (Thermo-Fisher Scientific) supplemented with 8% fetal bovine serum (Life Technologies), penicillin and streptomycin. KM12 (CMS1) was cultured in RPMI 1640 with L-glutamine, 25 mM HEPES (Thermo-Fisher Scientific) supplemented with 8% fetal bovine serum (Life Technologies), penicillin and streptomycin, 1% D-glucose solution plus (Sigma-Aldrich) and 100 μM sodium pyruvate (Thermo-Fisher Scientific). Co123 (CMS1) and Conc (CMS4) were cultured as spheroids in polyHEMA (poly(2-hydroxyethyl methacrylate), Sigma-Aldrich) coated flasks (Corning), in Advanced DMEM/F12 (Gibco) with 1:100 N2 (Invitrogen), 2 mM L-glutamine (Sigma-Aldrich), 5 mM HEPES (Life Technologies), 0.15% D-glucose (Sigma-Aldrich), 100 μM β-mercaptoethanol (Sigma-Aldrich), 10 μg/mL insulin (Sigma-Aldrich), 2 μg/mL heparin (Sigma-Aldrich), 1:1000 trace elements B and C (Corning), EGF 20 ng/mL and FGF 10 ng/mL. Hutu80, and KM12 were obtained from the Sanger Institute (Cambridge, UK), Co123 and Conc were established as described before^15^. All cell lines were authenticated by STR Genotyping and regularly tested for mycoplasma infection.

### Animal experiments

All *in vivo* experiments were approved by the animal experimentation committee of Amsterdam UMC under the nationally registered license number AVD118002016493 and performed according to national guidelines. NOD.Cg-Prkdc^scid^ Il2rg^tm1Wjl^ / Szj (NSG) mice were bred in-house.

### Establishment of HIS-NSG mice

Newborn NSG mice were sub-lethally irradiated once (1 Gy) using a 137Cs source and humanized by intrahepatic injection of human CD34^+^CD38^-^lineage^-^ cells (5 × 10^4^ cells) at five days postnatally. Eight weeks later, peripheral blood was collected from the submandibular vein to determine the reconstitution of a HIS which was deemed successful if the human immune cell engraftment score reached >20%. Mice were housed in individually ventilated cages with sterile bedding, food, and acidified water *ad libitum*.

### Subcutaneous tumor growth

To generate *in vivo* tumors, 50,000 CRC cells were resuspended in medium and mixed at a 1:1 ratio with Matrigel (Corning) and injected subcutaneously into both flanks of (HIS)-NSG mice. Tumor growth was measured twice a week using a caliper, using the formula 0.5 x length x width x height. Tumors from both flanks were pooled for each mouse prior to downstream analyses.

### Tissue dissociation and preparation for scRNA-seq

To isolate single cells for scRNA-seq, tumors were transferred to a 6-well plate and washed with ice cold PBS. Tumors were cut into small pieces using a scalpel blade and transferred to a 50 mL tube, and 10 mL warm (37 °C) digestion medium (RPMI 1640, 1.5 mg/mL collagenase, 20 µg/mL DNase I) was added. Samples were incubated at 37 °C for 30 min under continuous stirring. To stop the digestion process, 3 ml of cold wash medium (RPMI 1640 + 8% FCS) was added, the suspension was passed through a 70 µm cell strainer (Greiner) and spun down at 1500 rpm for 7 min. The pellet was resuspended in 2 ml red blood cell lysis buffer (eBioscience) and incubated for 5 min at RT. Eight ml of cold wash medium was added and the suspension was spun down at 1500 rpm for 7 min and washed twice and resuspended in FACS buffer (PBS + 0.1% BSA).

Spleens were pushed through a 100 µm cell strainer (Greiner) with a syringe plunger and spun down at 1500 rpm for 7 min. The pellet was resuspended in 5 ml red blood cell lysis buffer (eBioscience) and incubated for 5 min at RT. Ten ml FACS buffer (PBS + 0.1% BSA) was added, and the suspension was forced through 70 µm cell strainer (Greiner). After spinning at 1500 rpm for 7 min, the sample was resuspended in FACS buffer (PBS + 0.1% BSA). Tumor and spleen cells were counted and brought to a concentration of 25 x 10^6^ /mL for subsequent sorting.

### Flow cytometry

Cells were stained with anti-human CD45 APC (BD Biosciences) for 30 min at 4 ^̊^C and washed twice. Alive CD45^+^/DAPI^-^ cells were sorted using the SH800 Cell Sorter (Sony).

### scRNA-seq library preparation

Sorted anti-human CD45^+^/DAPI^-^ live cells were spun down at 493 x *g* for 5 min and supernatant was removed. Cells were diluted in 0.1% BSA/PBS to 1000 cells/μl and kept on ice until further processing. Directly after this, libraries were prepared according to Chromium Next GEM Single Cell 3ʹ Reagent Kits v3.1 (PN-1000121). Briefly, Gel Beads-in-emulsion (GEMs) were generated by combining barcoded Single Cell 3′ v3.1 Gel Beads, a Master Mix containing cells, and Partitioning Oil onto Chromium Next GEM Chip. Immediately following GEM generation, the Gel Bead was dissolved, primers were released, and any co-partitioned cell was lysed. Primers were mixed with the cell lysate and a mastermix containing reverse transcription (RT) reagents. Next, Silane magnetic beads were used to purify the first-strand cDNA from the post GEM-RT reaction mixture. The full-length cDNA was amplified via PCR to generate sufficient mass for library construction. After end repair, A-tailing, adaptor ligation and PCR amplification, the final libraries were ready for sequencing on the Illumina HiSeq 4000.

### Hashtags

Sorted anti-human CD45^+^/DAPI^-^ live cells were spun down at 493 x *g* for 5 min and supernatant was removed. Cells were transferred to an Eppendorf tube in 50 μl of 0.1% BSA/PBS and incubated with TotalSeq™ – B anti-human hashtag antibodies on ice for 30 minutes. After which cells were washed twice with 0.1% BSA/PBS and spun down at 360 x *g* for 5 minutes. Cells were counted and 5,000-10,000 cells per sample were used for pooling and kept on ice until further processing.

### scRNA-seq data analysis

Raw scRNA-seq reads were aligned to the combined human (GRCh38) and mouse (GRCm38) reference genome using the CellRanger 9.1.0 software from 10xGenomics to generate unique molecular identifier (UMI) counts. The filtered gene expression matrices were imported into R version 4.4 and further processed by the Seurat R package^16^ version 5.3.0 (filter cells with > 200 unique gene counts (nFeature RNA) and normalized using SCTransform^17^ using default parameters. Cellular quality control was conducted by calculating the percentage mitochondrial reads and cell cycle genes as proxies for cell death and proliferating cells. Cells were annotated using a semi-automatic approach, where automatic annotations were based on Seurat reference mapping against a human reference PBMCs CITE-seq experiment^18^, after which manual curation was performed based on canonical markers. For downstream analyses, human, non-proliferating, living immune cells were retained. Cell clusters were visualized with Uniform Manifold Approximation and Projection (UMAP) based on an initial selection of principal components (PCs). The PCs were chosen to represent those contributing 5% of the standard deviation individually and cumulatively accounting for 90% of the total standard deviation. The selection continued until the percentage change in variance between consecutive PCs dropped below 0.1%.

Publicly available raw scRNA-seq reads from mouse and human spleens were acquired from Hilton et al. 2019^19^ and the Tabula Sapiens Consortium et al. 2022^13^ as stored under Gene Expression Omnibus (GEO) accession numbers GSE132642 and GSE201333, respectively. Preprocessing of raw reads was performed as described above. Publicly available processed scRNA-seq data in the form of UMI count matrices were obtained from human tumors reported by Lee et al., available under GEO accession GSE132465. The preprocessing of these data followed the same procedure outlined previously. Cell-type annotations were retrieved from the original publications and harmonized with those used in our dataset.

### Immunohistochemistry

Directly after isolation, tumors were fixed in 4% paraformaldehyde overnight prior to paraffin embedding. Tissue sections (5 μm) were deparaffinized and antigen retrieval was performed using 10 mM sodium citrate and boiling for 20 min. Endogenous peroxidase activity was blocked with 3% hydrogen peroxide in PBS. Non-specific staining was blocked using UltraVision Protein Blk (Thermo Scientific, Waltham, MA) 10 min on RT. Primary antibodies CD4 (Abcam, 1:50), CD8 (DAKO, 1:50), CD20 (DAKO, 1:500), CD56 (Monosan, 1:50), CD163 (Cell Marque Corporation, 1:25) were diluted in antibody diluent (Agilent: CD4; Ventana: CD8, CD20, CD56, CD163) and incubated in a humidified chamber. For amplification of the staining, Brightvision+ post antibody block (Immunologic) was used for 20 min prior to the addition of the secondary antibody, poly-HRP-anti Ms/Rb IgG (Immunologic) for 30 min at RT. Visualization of stainings was performed with Bright DAB solution (Immunologic) according to manufacturer’s protocol, counterstained with undiluted Mayer Haematoxylin (Klinipath) and mounted tissue sections with non-aqueous medium.

### Statistics

Single-cell differential abundance analyses were conducted using integrated statistical analysis using GraphPad (Mann-Whitney tests). Differential expression analyses were conducted using the pseudobulk method^20^ by aggregating reads per cell type per sample and using DESeq2 for statistical testing^21^. Gene set enrichment analyses were conducted using the fgsea package.

## RESULTS

### Systemic immune landscape in HIS mice defined by single-cell RNA sequencing

To characterize the systemic immune landscape in HIS mice, spleens from HIS mice without tumors (control; n = 6) were processed into single-cell suspensions, and scRNA-seq was performed on FACS-sorted human CD45^+^ immune cells to profile their transcriptional states (fig 1A and Supplementary Table 1).

**Figure 1.**
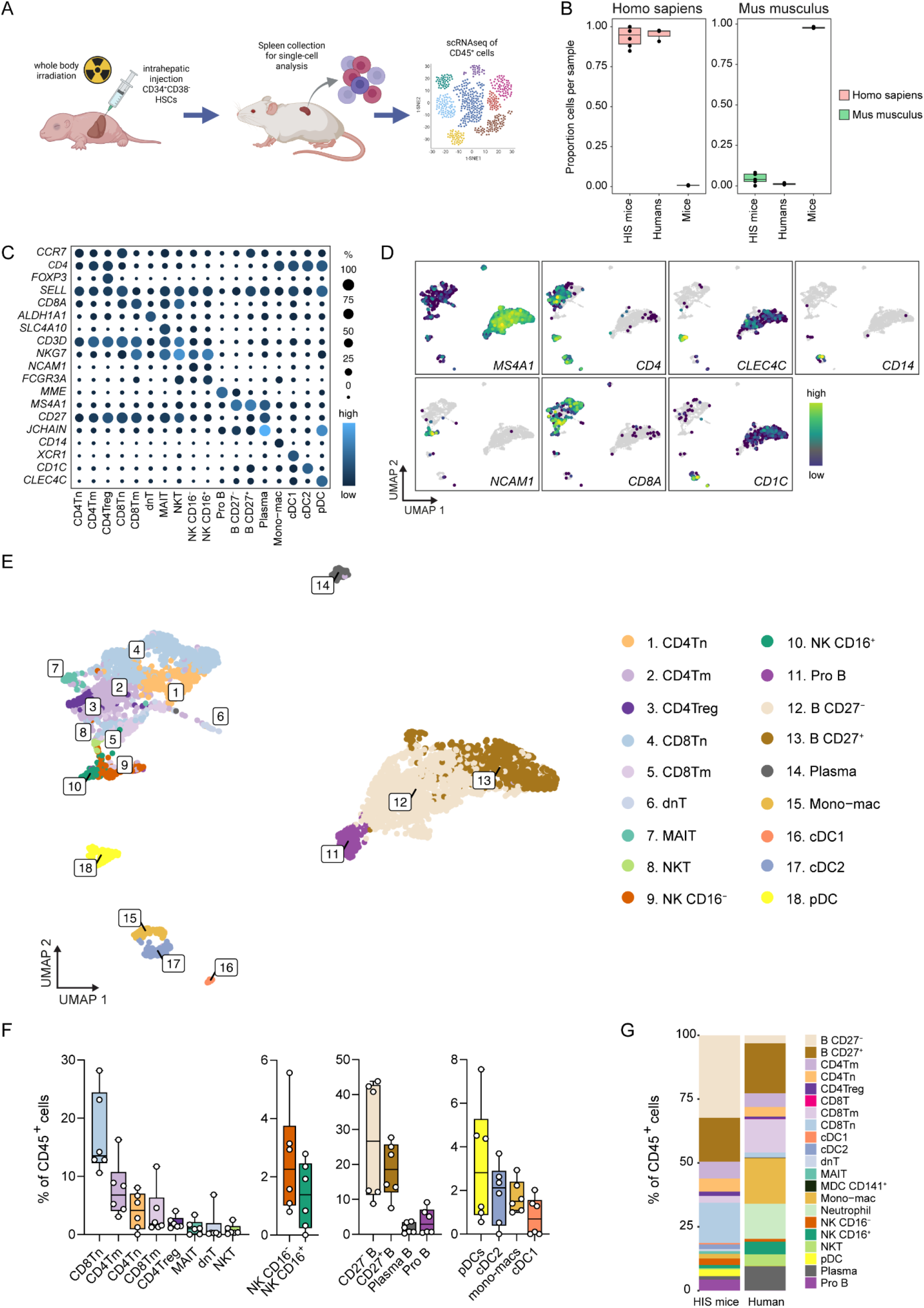
Immune landscape in HIS mice defined by single-cell RNA sequencing. **(A)** Schematic overview of part of the HIS mouse experiment. Spleens from HIS mice without tumor implantation (control; n = 6) were collected, dissociated and analyzed by scRNA-seq. Created in BioRender. Grootjans, J. (2026) https://BioRender.com/9mopybk **(B)** Boxplot showing the proportion of splenic cells with >90% of transcriptomic read mapping to either the *Homo sapiens* (GRCh38) or the *Mus musculus* (GRCm38) reference genome in control HIS mice (n = 6), human spleens^13^ (n = 6), and mouse spleens^19^ (n = 4). **(C)** Dotplot of canonical marker gene expression of different immune subsets, where dot size represents the proportion of cells expressing the gene, and color indicates expression level. **(D)** UMAP feature plots showing log-transformed expression of canonical marker genes *MS4A1, CD4, CLEC4C, CD14, NCAM1, CD8A, and CD1C*. **(E)** UMAP colored by immune cell subset annotation (n = 5,633 cells). **(F)** Boxplot showing proportion of immune cell subsets relative to total CD45^+^ splenocytes in HIS mice (n = 6). **(G)** Graph showing immune cell subset proportions relative to total CD45^+^ splenocytes in HIS mice (n = 6) and human spleens (n = 3). B: B cell; T: T cell; Tn: T naive; Tm: T memory; Treg: regulatory T; MAIT: mucosal associated invariant T; NKT: natural killer T; dnT: double negative T; NK: natural killer cell; pDC: plasmacytoid dendritic cell; mono-macs: monocytes-macrophages; cDC1: conventional dendritic cell type 1; cDC2: conventional dendritic cell type 2; UMAP: Uniform Manifold Approximation and Projection; HIS: humanized immune system.

To confirm that the immune cells originated from humans, we mapped the scRNA-seq reads to a combined human (GRCh38) and mouse (GRCm38) reference genome and compared the resulting profiles to publicly available splenic scRNA-seq datasets from both species. The majority of reads aligned to the human genome rather than the murine one, more closely resembling human than murine scRNA-seq data (fig 1B). For subsequent analyses, cells mapping to the murine genome were discarded. Based on canonical marker gene expression, we annotated 18 distinct immune cell subsets (fig 1C,D). These subsets were further characterized using marker gene analyses to identify differentially expressed genes (DEGs) (fig S1A). Immune cell populations were consistently detected across individual HIS mice and donor mixes, with no immune subsets uniquely associated with a specific mouse or donor source (fig S1B,C).

Using Uniform Manifold Approximation and Projection (UMAP) analysis of all included CD45^+^ cells (n = 5,633 cells), we visualized and annotated the systemic human immune cell landscape present in HIS mice (fig 1E). For comparison purposes, immune populations were named according to established human immune definitions, and all major immune lineages typically present in humans were identified. We quantified a total of eight T cell subsets, two natural killer (NK) cell subsets, four B cell subsets, and four myeloid subsets in the spleens of HIS mice.

Among human CD45^+^ immune cells in HIS mouse spleens, a large fraction consisted of CD27^-^ B cells (mean: 26.7%) and CD27^+^ B cells (mean: 18.4%), as well as Plasma B cells (mean: 3.8%) and pro B cells (mean: 1.8%). Within the T cell lineage, the most abundant subset was naive CD8^+^ T cells (CD8Tn; mean: 16.1%), followed by memory CD4^+^ T cells (CD4Tm; mean: 7.7%), naive CD4^+^ T cells (CD4Tn; mean: 4.0%), memory CD8^+^ T cells (CD8Tm; mean: 3.7%), regulatory CD4^+^ T cells (CD4Treg; mean: 2.0%), mucosal-associated invariant T cells (MAIT; mean: 1.2%), double-negative T cells (dnT; mean: 1.7%), and natural killer T cells (NKT; mean: 0.8%).

The NK cells comprised two populations, NK CD16^-^ (mean: 2.5%) and NK CD16^+^ cells (mean: 1.4%). Finally, four major myeloid populations were identified, in decreasing order of abundance: plasmacytoid dendritic cells (pDCs; mean: 3.3%), conventional DCs type 2 (cDC2; mean: 1.8%), monocyte-macrophages (mean: 1.6%) and conventional DCs type 1 (cDC1; mean: 0.8%) (fig 1F).

To benchmark these findings against the human systemic immune landscape, we compared our data with a publicly available scRNA-seq dataset of healthy human spleens (n = 3) provided by Tabula Sapiens^13^. This comparison confirmed that all major immune cell types present in human spleen were also detected in HIS mice (fig 1G). Notably, CD141^+^ myeloid DCs (MDC) and neutrophils were detected in human spleen samples but were not observed in spleens from HIS mice (fig 1G).

Together, these data demonstrate that the systemic immune compartment of HIS mice can be robustly characterized by scRNA-seq and broadly reflects the cellular composition of the human immune system, supporting the suitability of this model for studying systemic human immune responses *in vivo*.

### Human-like immune infiltration in subcutaneously grown human CRC in HIS mice

To investigate the immune composition of human CRC tumors in HIS mice, four cell lines (CMS1: KM12 and Co123; CMS4: Hutu80 and Conc) were subcutaneously injected in mice. All cell lines formed tumors successfully (fig S2A). To characterize the tumor-associated immune compartment, we performed scRNA-seq on human CD45^+^ immune cells isolated from 11 mice, yielding a total of 14,578 cells. Tumor-derived immune cells were annotated based on canonical marker gene expression (fig 2A,B), supported by differential gene expression analyses (fig S2B). All 18 human CD45^+^ immune subsets identified in the spleen were also detected in tumors (fig 2C).

**Figure 2.**
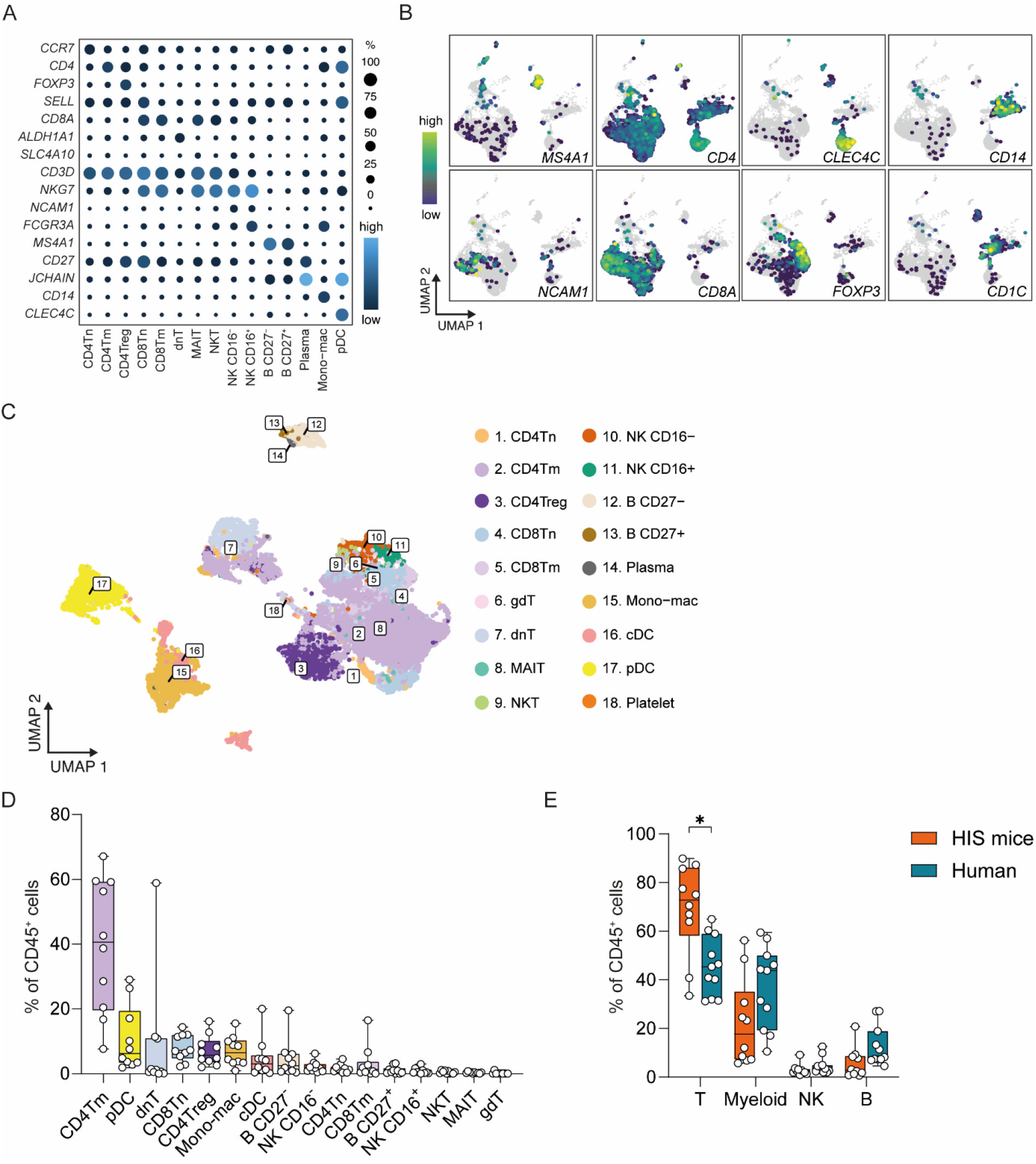
Human immune cell infiltration in subcutaneous grown human CRC in HIS mice. **(A)** Dotplot of canonical marker gene expression of different immune subsets in tumors of HIS mice (n = 10), where dot size represents the proportion of cells expressing the gene and color indicates expression level. **(B)** UMAP feature plots of CD45^+^ immune cells from HIS mouse tumors showing log-transformed expression of canonical marker genes *MS4A1, CD4, CLEC4C, CD14, NCAM1, CD8A, FOXP3 and CD1C*. **(C)** UMAP colored by immune cell subset annotation in CD45^+^ immune cells from HIS mouse tumors (n = 14,578 cells). **(D)** Boxplot showing the proportion of immune cell subsets relative to total CD45^+^ immune cells in HIS mouse tumors (n = 10 mice), ordered by mean abundance. **(E)** Boxplot comparing proportions of major immune lineages relative to CD45^+^ immune cells in HIS mouse tumors (n = 10) and human CRC tumors (n = 11). Statistics: Mann-Whitney test, two-tailed. Only statistically significant differences are shown (*p ≤ 0.05). B: B cell; T: T cell; Tn: T naive; Tm: T memory; Treg: regulatory T; CD4Tfh: CD4 T follicular helper cell; CD4Th17: CD4 Th17 cell; MAIT: mucosal associated invariant T; NKT: natural killer T; dnT: double negative T; NK: natural killer cell; pDC: plasmacytoid dendritic cell; mono-macs: monocytes-macrophages; cDC1: conventional dendritic cell type 1; cDC2: conventional dendritic cell type 2; UMAP: Uniform Manifold Approximation and Projection; HIS: humanized immune system. NSG: NOD scid gamma; CMS: Consensus Molecular Subtype.

Among tumor-infiltrating immune populations, CD4Tm cells were the most abundant immune subset (mean 39.7%). Other T cell subsets infiltrating tumors were mainly dnT cells (mean 8.6%), CD8Tn cells (mean 8.1%) and CD4Treg cells (mean 7.1%). Myeloid and innate tumor-infiltrating immune populations were particularly pDCs (mean 10.7%) and monocyte-macrophages (mean 7.0%) (fig 2D).

To evaluate to what extent the immune infiltrate in HIS mouse tumors reflects that of human CRC, we compared our data with immune cells derived from 11 primary human CRC tumors (CMS1 n=5, CMS4 n=6) in a publicly available scRNA-seq dataset (GSE132465)^14^ (n = 13,451 cells). At the level of major immune lineages, HIS mouse tumors contained a significantly higher proportion of T cells compared with human tumors (mean 69.1% vs. 45.6%). Although not statistically significant, myeloid cells (mean 22.4% vs. 37.0%) and B cells (mean 5.6% vs. 12.7%) were less abundant in HIS mouse tumors compared to human tumors. No substantial difference was observed in the overall NK cell fraction between HIS and human tumors (mean 2.9% vs 4.7%) (fig 2E).

Further analysis revealed a significantly increased CD4/CD8 T cell ratio in HIS mouse tumors compared with human CRC tumors (fig S2C). As expected, more pronounced differences emerged when examining individual immune subpopulations (fig S2D). Within the T cell compartment, CD4^+^ effector memory T cells were markedly overrepresented in HIS tumors compared with human tumors (56.3% vs. <1%), whereas CD4^+^ regulatory T cells were present in both, albeit at lower frequencies in HIS mice (10.0% vs. 25.3%). Within the myeloid compartment, HIS mouse tumors contained fewer monocyte-macrophages relative to their parent lineage (mean 36.8% vs. 89.7% of myeloids) and relatively higher proportions of pDCs (mean 46.1% vs. 3.3%) and cDCs (mean 17.1% vs. 5.1%) compared with human tumors. In contrast, NK cell subsets showed the closest correspondence between HIS mouse and human tumors, with comparable proportions of CD16^−^ NK cells (mean 70.0% vs. 66.8%) and CD16^+^ NK cells (mean 30.1% vs. 33.2%), respectively (fig S2D). Within the B cell compartment, CD27^−^ B cells represented the dominant population in HIS mouse tumors, whereas this was less prevalent in human tumors (mean 76.7% vs. 8.2%). Several immune cell subsets in the human CRC dataset^14^ were not identified in our HIS mouse tumors, including plasma cells, proliferating myeloid cells, CD4^+^ follicular helper T cells (CD4Tfh) and CD4^+^ Th17 cells.

Collectively, these results demonstrate that human CRC tumors grown in HIS mice develop a complex immune infiltrate resembling human tumors, encompassing multiple immunologically relevant cell populations. Although differences in immune composition remain, the presence of key T cell, myeloid, and NK-cell subsets supports the potential utility of this model for preclinical immuno-oncology research.

### Transcriptional profile of CMS-dependent immune infiltration recapitulated in HIS mice

We next investigated whether immune infiltration in subcutaneous CRC tumors grown in HIS mice differed according to the consensus molecular subtype (CMS) of the injected cell lines. At the level of overall immune composition, no major differences in immune cell abundance were observed between CMS subtypes (fig 3A). Quantitative analysis confirmed that no statistically significant differences in immune cell proportions were detected across CMS groups (fig 3B).

**Figure 3.**
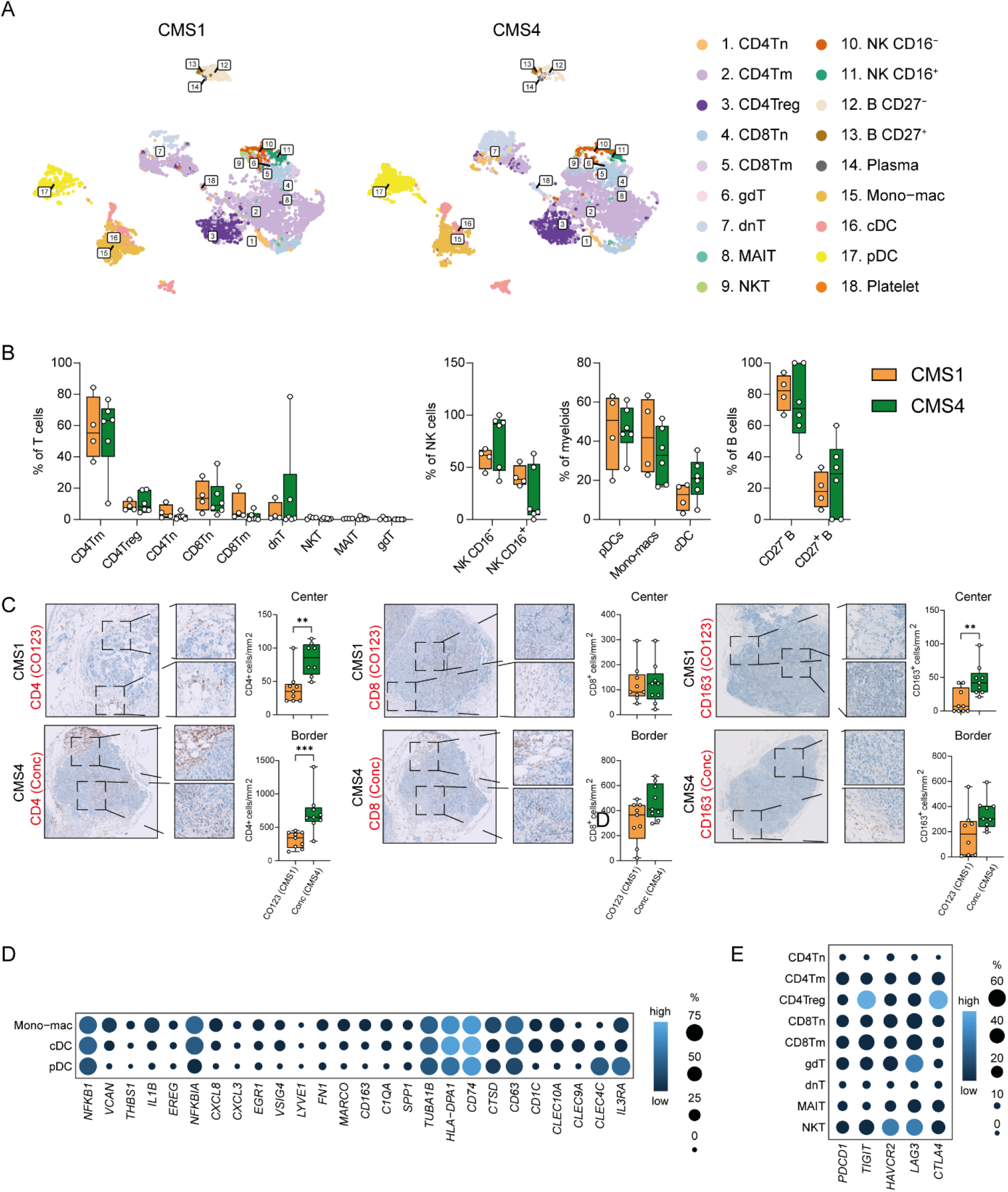
T cell exhaustion recapitulated in HIS mice. **(A)** UMAP of CD45^+^ immune cells from HIS mouse tumors, split per CMS subtype (n = 14,578 cells). **(B)** Boxplots showing proportions of tumor-infiltrating immune cells across major lineages (T, NK, myeloid, and B cells) stratified by CMS subtype. **(C)** Representative immunohistochemical staining of FFPE sections from subcutaneous HIS CRC tumors of CMS1 and CMS4 subtypes using CD4, CD8, and CD163, with corresponding boxplot quantification of CD4^+^, CD8^+^, and CD163^+^ cells expressed as cells/mm^2^. Statistical significance was determined using a two-tailed Mann–Whitney test. Only statistically significant differences are shown (**p ≤ 0.01; ***p ≤ 0.001). **(D)** Dotplot of genes associated with human macrophage characterization^22^ across myeloid subsets in HIS tumors. **(E)** Dotplot of genes associated with human T cell exhaustion across T cell subsets in HIS tumors. Dot size indicates the proportion of cells expressing the gene, and color indicates expression level. B: B cell; T: T cell; Tn: T naive; Tm: T memory; Treg: regulatory T; CD4Tfh: CD4^+^ follicular helper T cell; CD4Th17: CD4^+^ Th17 cell; MAIT: mucosal associated invariant T; NKT: natural killer T; dnT: double negative T; NK: natural killer cell; pDC: plasmacytoid dendritic cell; mono-macs: monocytes-macrophages; cDC1: conventional dendritic cell type 1; cDC2: conventional dendritic cell type 2; UMAP: Uniform Manifold Approximation and Projection; HIS: humanized immune system; CMS: Consensus Molecular Subtype.

Despite the absence of marked compositional differences by scRNA-seq, immunohistochemical analyses revealed CMS-associated spatial differences in immune cell distribution. CMS4 tumors exhibited a relatively higher abundance of CD4^+^ T cells in both the tumor center and invasive margin, as well as increased infiltration of CD163^+^ monocyte-macrophages in the tumor center (fig 3C). In addition, quantification of NK cells and B cells using CD56 and CD20 staining, respectively, demonstrated an increased presence of NK cells at the tumor border of CMS4 tumors (fig S3A). To determine whether transcriptional programs of tumor-associated myeloid populations reflected human-like immune features, we examined expression of a curated gene set comprising established macrophage markers and canonical DC genes^22^. All myeloid subsets displayed high expression of genes including *NFKB1, NFKBIA, TUBA1B, HLA-DPA1* and *CD74* (fig 3D), which are associated with both pro- and anti-inflammatory immune functions^23-27^. Furthermore, using a predefined gene set^28,29^ associated with wound-healing programs, a hallmark of myeloid cells in cancer^30,31^, these populations were characterized by elevated expression of *CNN2, RAF1, ELK3, TGFBR1, TGFBR2, CD44, PAK1* and *MACF1* (fig S3B). The majority of these genes are linked to anti-inflammatory and pro-tumoral macrophage phenotypes consistent with prior observations in human CRC^22,32-34^(fig S3B). Given the emerging role of ICB in CRC, we next examined the expression of genes associated with T cell exhaustion^35^. In HIS mouse tumors, multiple T cell subsets showed high expression of *PDCD1, TIGIT, HAVCR2, LAG3* and *CTLA4*. Naïve CD4^+^ T cells exhibited minimal expression of these genes, consistent with their differentiation state, whereas CD4^+^ regulatory T cells displayed particularly high levels of *TIGIT* and *CTLA4* (fig 3E).

Collectively, these data indicate that, despite limited numerical differences in immune cell composition across CMS subtypes, HIS mouse tumors recapitulate key transcriptional programs associated with immune regulation and T cell dysfunction. This supports the utility of HIS mice as a biologically relevant platform for investigating the tumor immune microenvironment in CRC and for the preclinical evaluation of immune checkpoint–based therapies.

## DISCUSSION

In this study, we provide a comprehensive immunological characterization of both the systemic immune system and the TIME in a CRC HIS mouse model using scRNA-seq. By benchmarking splenic and tumor-associated immune compartments against human datasets, we were able to evaluate the extent to which this model recapitulates human immune cell composition and transcriptional states at single-cell resolution. Based on gene expression profiles, we identified all major immune subsets in the spleen of HIS mice, and their relative abundance closely resembled previously published HIS mouse studies in which splenic immune cells were phenotyped using flow cytometry^36^. The high abundance of immature pro B cells observed in HIS mouse spleens is consistent with known limitations of these models, as B cell maturation and differentiation are impaired^37^.

Comparison of tumor-infiltrating immune cells from HIS mouse tumors with those from primary human CRCs revealed the presence of largely overlapping immune cell subsets, indicating that key elements of the human CRC immune landscape are captured in this model. In line with previous reports, we observed a predominance of CD4^+^ memory T cells, which has been suggested to arise as a compensatory mechanism following lymphopenia during immune reconstitution in HIS mice^38^. Moreover, differences in cell-type annotation strategies between our dataset and the reference study may partially account for this discrepancy. In addition, the relatively high proportion of pDCs is consistent with earlier reports^39^. Compared with human CRC tumors, HIS mouse tumors displayed a higher overall fraction of T cells and a relative underrepresentation of myeloid cells, likely as a result of defective myeloid immune cell reconstitution in HIS mice^40^. Nevertheless, the presence of a substantial myeloid compartment (mean ∼22%) suggests that HIS mice still enable meaningful investigation of myeloid-T cell interactions within the tumor microenvironment.

Although scRNA-seq did not reveal major proportional differences in immune cell subsets across CMS subtypes, immunohistochemical analyses demonstrated CMS-associated spatial differences in immune infiltration. Specifically, CMS4 tumors showed increased infiltration of CD4^+^ T cells and CD163^+^ monocyte-macrophages in both the tumor center and invasive margin. These findings suggest that spatial organization and local enrichment of immune cells, rather than global immune abundance alone, may better capture CMS-associated immune features in this model.

Interestingly, myeloid cells in HIS mouse tumors displayed transcriptional programs consistent with anti-inflammatory and wound healing signatures, which are known to contribute to tumor progression^41^. However, we did not observe preferential enrichment of these programs in the CMS4 subtype, as reported in human CRC^9^. This discrepancy may reflect limitations of the subcutaneous tumor model, the absence of stromal components of human origin, or limited myeloid lineage maturation in HIS mice. Alternative humanized mouse models with improved myeloid lineage reconstitution have been described and may be even more suitable for studies focusing on myeloid-driven tumor phenotypes, such as CMS4 CRC^42^. However, to further assess the functional state of myeloid cells present, we examined the expression of genes previously associated with macrophage activation and dendritic cell identity and observed broad expression of genes such as *NFKB1* and *HLA-DPA1*. The concurrent expression of genes linked to both pro- and anti-inflammatory functions underscores the plastic and dynamic nature of tumor-associated myeloid cells within the HIS mouse microenvironment^23-27^.

To further uncover T cell biology in light of recent clinical breakthroughs^35,43,44^, we performed an in-depth analysis of T cell states within HIS mouse tumors. We identified nine distinct human T cell subsets that closely correspond to those described in primary human CRC tumors. Notably, exhaustion-associated genes, including *PDCD1, TIGIT, HAVCR2, LAG3* and *CTLA4*, were expressed across multiple T cell subsets, whereas naïve CD4^+^ T cells showed minimal expression of these markers, consistent with their differentiation state. In line with our data, T cell exhaustion (i.e. PD-1, TIGIT and CD27) was previously observed in splenic T cells from HIS mice with PDX tumors from breast cancer patients^36^. CD4^+^ regulatory T cells exhibited particularly high expression of *TIGIT* and *CTLA4*. This finding is of interest in light of ongoing clinical development of TIGIT-targeted therapies aimed at restoring anti-tumor T cell function^45-48^. Furthermore, the widespread expression of *PDCD1* supports the translational relevance of this model and aligns with recent studies demonstrating therapeutic efficacy of PD-1-based combination therapies in HIS mouse models of MSS CRC^8^.

This study has several limitations. First, neutrophils were present at low abundance in HIS mice, which may reflect technical constraints of the scRNA-seq platform used^49^. Second, despite stringent filtering for human-aligned reads, a minor contribution of murine stromal or immune components cannot be fully excluded and may influence immune cell behavior within the tumor microenvironment. Finally, although the CRC cell lines used in this study were not HLA-matched to the HSC donors, previous studies have shown that CD34^+^ HSC-based HIS mice lack fully HLA-restricted T cell education due to the murine thymic environment, thereby limiting the impact of HLA mismatch in this setting^50^.

In conclusion, our study demonstrates that HIS mice recapitulate key aspects of both the systemic immune compartment and the CRC tumor immune microenvironment at single-cell resolution. By directly benchmarking immune cell composition and transcriptional programs against human datasets, we provide a framework to assess the translational relevance of HIS mouse models. Importantly, the presence of human-like T cell exhaustion states supports the use of HIS mice as a preclinical platform for investigating immunomodulatory strategies, including immune checkpoint blockade, in MSS CRC. Together, these findings highlight both the strengths and limitations of HIS mouse models and provide guidance for their rational application in immuno-oncology research.

## Data availability statement

All raw sequencing data generated during this study have been deposited in the European Genome-phenome Archive (EGA) under controlled access to comply with the local privacy laws. All data can be found on Zenodo, DOI: 10.5281/zenodo.18754883. Researchers are allowed to apply for access to the data access committee governing this data under the condition that the data will be used for research purposes only. Data from human spleen samples were obtained from GSE201333. Data from human CRC tumors were obtained from GSE132465.

## Code availability

All computational scripts used to prepare and analyze the data can be found on https://github.com/AmsterdamUMC/ELNID001000000000472986

## Acknowledgements

We wish to thank the staff from all the units that participated in the study. In particular, Susan Kenter, Core facility Genomics and Lisanne Nijman, support with laboratory experiments. Toni van Capel, Berend Hooibrink, Kim Brandwijk-Paarlberg and Daisy Picavet-Havik, Core facility Advanced Microscopy and Flow Cytometry and Wim Vos, Pathology. Fig 1A was created with BioRender.com. This work is supported by The New York Stem Cell Foundation (LV), grants from the European Research Council (ERC-CoG 101045612 - NIMICRY) (LV), ZonMw (VICB 09-15018-21-10029) (LV) and Dutch Cancer Society grant (KWF 13435 / 2021-1) (LV), KWF YIG 13915 (JG), NWO ZonMw Veni Grant 09150161810115 (JG), NWO ZonMw Vidi Grant 09150172210058 (JG), Top Institute for Knowledge and Innovation grant ImPACT (JG), donation by mr H.J.M. Roels through Oncode Institute (JG).

## Author Contributions Statement

J.S., S.B. and F.A.V.B. performed experiments. J.S., S.B. and F.A.V.B. collected material. J.S., S.B., A.L.Y. and F.A.V.B. analyzed data. A.L.Y. provided bioinformatics input. J.S., S.B., J.G. and L.V. conceived, designed and directed the research. J.S., S.B., A.L.Y., J.G. and L.V. provided scientific input. J.S., A.L.Y. and S.B. wrote the manuscript. J.G. and L.V. supervised the manuscript. All authors approved the content of the manuscript.

## Supplementary Material

**Supplementary Figure 1.**
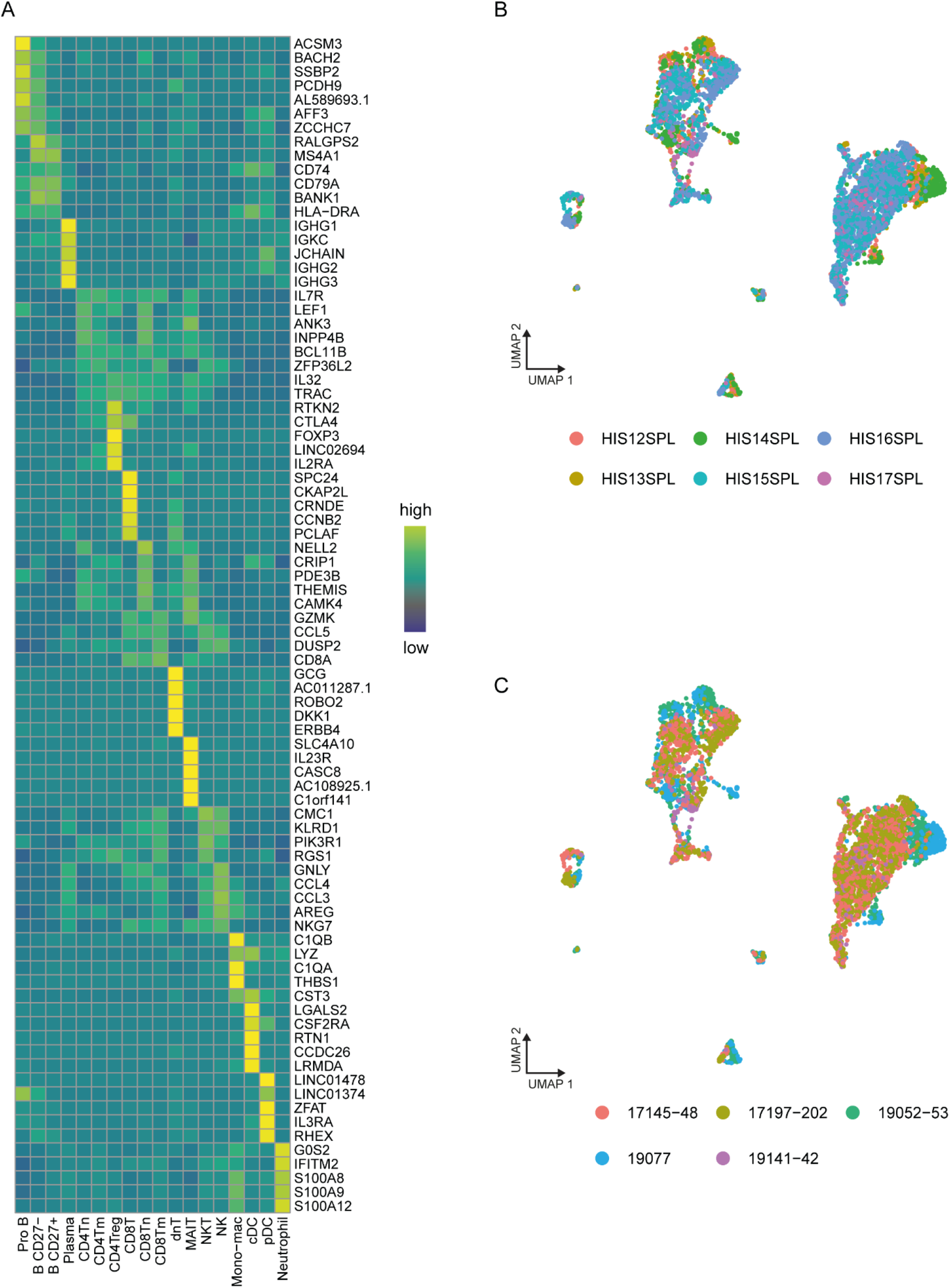
Systemic immune landscape in HIS mice defined by single-cell RNA sequencing. **(A)** Heatmap showing expression of the top five differentially expressed marker genes for each immune cluster identified among CD45^+^ splenocytes from HIS mice by scRNA-seq. **(B)** UMAP projection of all CD45^+^ splenocytes (n = 5,633 cells), colored by immune cell subtype and stratified by individual HIS mice. **(C)** UMAP projection colored by donor mix used to generate HIS mice. B, B cell; T, T cell; Tn, naïve T cell; Tm, memory T cell; Treg, regulatory T cell; MAIT, mucosal-associated invariant T cell; NKT, natural killer T cell; dnT, double-negative T cell; NK, natural killer cell; pDC, plasmacytoid dendritic cell; mono-macs, monocytes-macrophages; cDC1, conventional dendritic cell type 1; cDC2, conventional dendritic cell type 2; UMAP, Uniform Manifold Approximation and Projection.

**Supplementary Figure 2.**
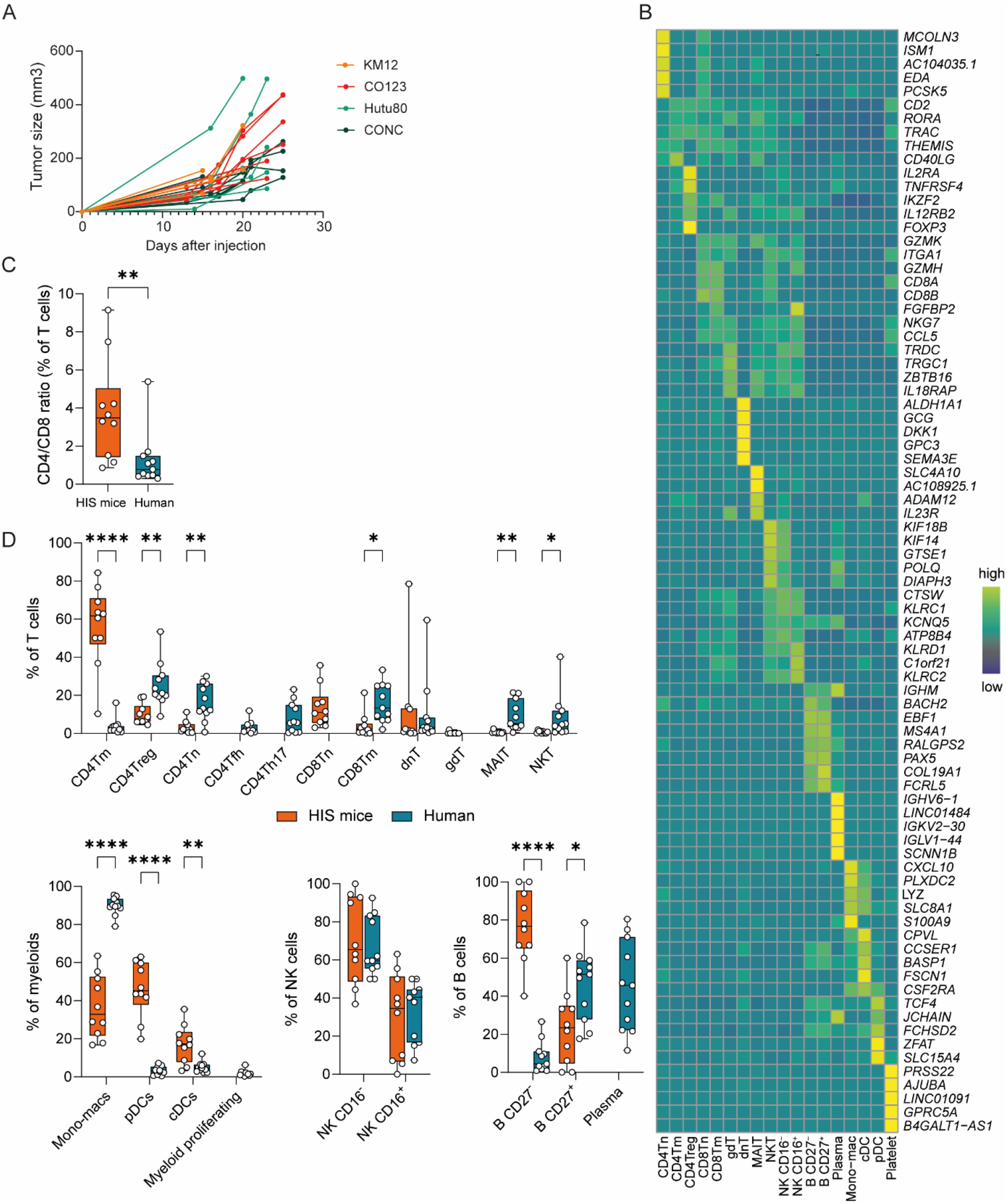
Human immune cell infiltration in subcutaneous grown human CRC tumors in HIS mice. **(A)** Tumor growth over time of KM12 (CMS1) CO123 (CMS1), Hutu80 (CMS4) and Conc (CMS4) CRC cell lines in NSG HIS mice. **(B)** Heatmap showing expression of the top five differentially expressed marker genes for each immune cluster identified among CD45^+^ immune cells isolated from subcutaneous CRC tumors in HIS mice by scRNA-seq. **(C)** Boxplot comparing CD4/CD8 T cell ratios in HIS mouse tumors and primary human CRC tumors, calculated from total T cell populations. **(D)** Boxplot showing the relative proportions of immune cell subsets within their respective major lineages in HIS mouse tumors (n = 10) and human CRC tumors (n = 11). Statistical significance was determined using a two-tailed Mann–Whitney test. Only statistically significant differences are shown (*p ≤ 0.05; **p ≤ 0.01; ***p ≤ 0.001; ****p ≤ 0.0001). B: B cell; T: T cell; Tn: naïve T cell; Tm: memory T cell; Treg: regulatory T cell; CD4Tfh: CD4 follicular helper T cell; CD4Th17: CD4 Th17 cell; MAIT: mucosal-associated invariant T cell; NKT: natural killer T cell; dnT: double-negative T cell; NK: natural killer cell; pDC: plasmacytoid dendritic cell; mono-macs: monocytes-macrophages; cDC1: conventional dendritic cell type 1; cDC2: conventional dendritic cell type 2; UMAP: Uniform Manifold Approximation and Projection; HIS: humanized immune system.

**Supplementary Figure 3.**
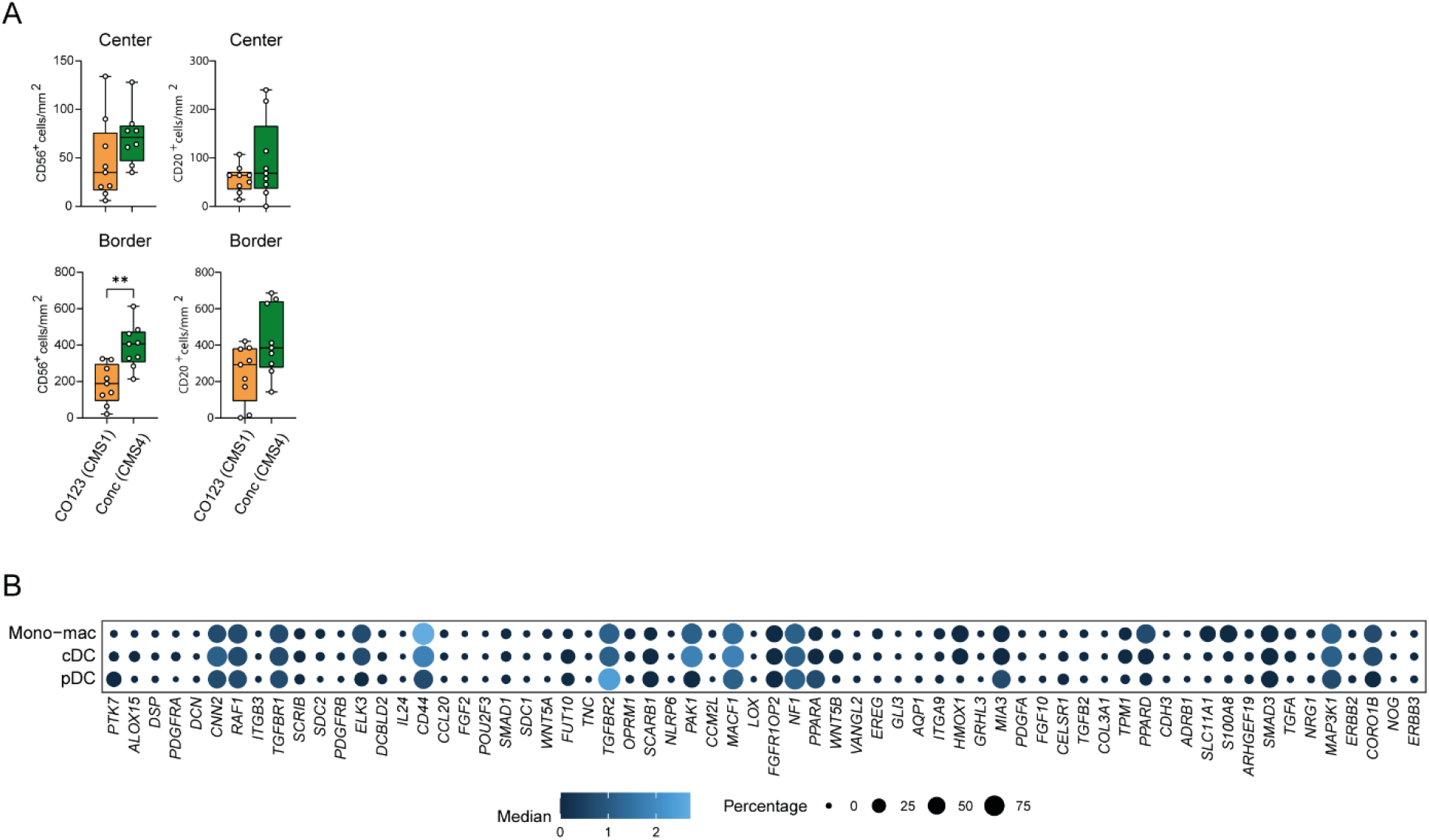
Additional characterization of tumor immune features in HIS mice. **(A)** Boxplot quantification of CD56^+^ NK cells and CD20^+^ B cells (cells/mm^2^) comparing tumor center and invasive margin (border) across different CMS subtypes. Only statistically significant differences are shown (**p ≤ 0.01). **(B)** Dotplot showing expression of wound-healing–associated genes across myeloid cell subsets in HIS tumors, where dot size indicates the proportion of cells expressing each gene and color intensity reflects expression level. Center: tumor center; Border: tumor border; CMS: Consensus Molecular Subtype; pDC: plasmacytoid dendritic cell; mono-macs: monocytes-macrophages; cDC: conventional dendritic cell.

